# Diterpenoid Vinigrol activates ATF4/DDIT3-mediated PERK/eIF2α arm of unfolded protein response to drive breast cancer cell death

**DOI:** 10.1101/2021.08.25.457587

**Authors:** Wencheng Wei, Yunfei Li, Chuanxi Wang, Sanxing Gao, Hao Wang, Yan Zhao, Ziying Gao, Yanxiang Jiang, Hao Gao, Xinsheng Yao, Yuhui Hu

## Abstract

Vinigrol is a natural diterpenoid with unprecedented chemical structure, driving great efforts into its total synthesis and the chemical analogs in the past decades. Despite its pharmacological efficacies reported on anti-hypertension and anti-clot, comprehensive functional investigations on Vinigrol and the underlying molecular mechanisms are entirely missing. In this study, we carried out a complete functional prediction of Vinigrol using a transcriptome-based strategy, Connectivity Map, and identified “anti-cancer” as the most prominent biofunction ahead of anti-hypertension and anti-depression/psychosis. A broad cytotoxicity was subsequently confirmed on multiple cancer types. Further mechanistic investigation on MCF7 cells revealed that its anti-cancer effect is mainly through activating PERK/eIF2α arm of unfolded protein response (UPR) and subsequent upregulation of p53/p21 to halt the cell cycle. The other two branches of UPR, IRE1α and ATF6, are functionally irrelevant to Vinigrol-induced cell death. CRISPR/Cas9-based gene activation, repression, and knockout systems identified essential contribution of ATF4/DDIT3 not ATF6 to the death process. This study unraveled a broad anti-cancer function of Vinigrol and its underlying targets and regulatory mechanisms, and also paved the way for further inspection on the structure-efficacy relationship of the whole compound family, making them a novel cluster of chemical hits for cancer therapy.

## 1. Introduction

Vinigrol is a diterpenoid with a brand new chemical structure originally purified from the fungal strain *Virgaria nigra* in 1987 ^1^. It is the only terpenoid consisting of two cis-fused ring systems and an eight-membered ring bridge, and thus persisted as a formidable challenge for chemical synthesis ^1-6^ (Fig. 1A). Large efforts from chemists in the past half century have led to a success in its total synthesis ^2-3^, which in turn raised questions for biologists on the potential medical applications of Vinigrol and its synthetic analogues. Unfortunately, the existing knowledge on Vinigrol’s bioactivities was solely based on one rudimentary study from early years, offering only the pharmacological observations on anti-hypertension and anti-platelet aggregation effects ^7,8^. Neither the underlying mechanisms nor the molecular targets of Vinigrol were investigated. Thus, a comprehensive and unbiased exploration on the functions and regulatory mechanisms of Vinigrol is pivotal to transforming the chemical success into the medical uses.

**Figure 1.**
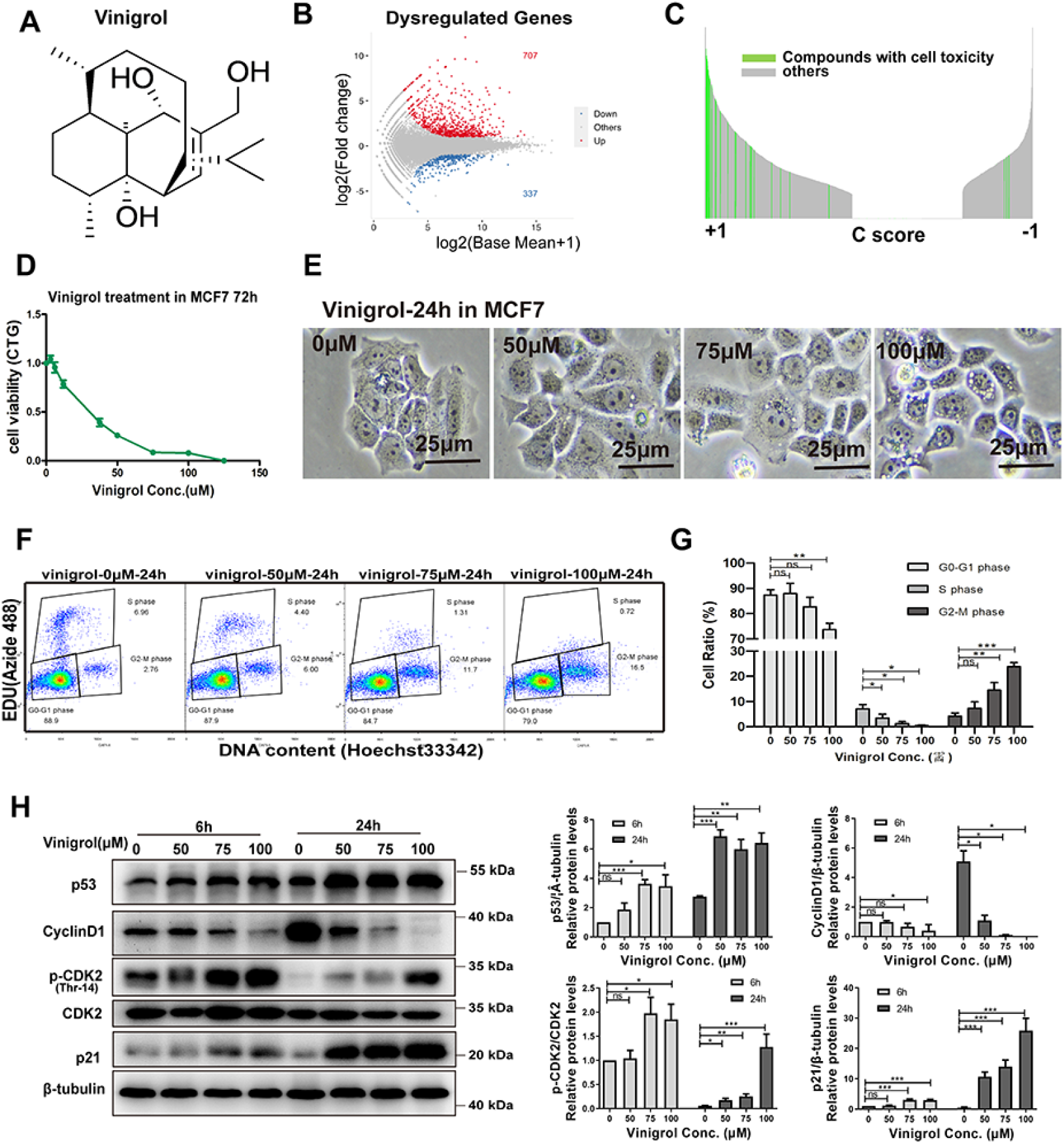
Vinigrol shows anticancer potency via DNA damage and cell cycle arrest. (A) Chemical structure of Vinigrol. (B) MA plot figure analysis by 3’RNA-seq. (C) Ranking of 6100 cScores of Vinigrol with the compounds in CMap Build02. The length of each line indicates the absolute value of cScore. Green lines represent toxicity compounds and grey lines represent non-toxicity compounds. (D) Cell viability after Vinigrol treatment 72 hrs measured by CTG assay. (E) Cell morphology after Vinigrol treatment 24 hrs. (F) Flow cytometry analysis of MCF7 cells treated 24 hrs for different Vinigrol doses, stained with EdU and Hoeschst, (G) Proportion of cells at different cell cycles after Vinigrol treatment 24 hrs at varied doses. (H) (Left) Western blot analysis of cell cycle checkpoint-related proteins after varied doses of Vinigrol treatment for 6 hrs and 24 hrs. (Right) semi-quantification of protein levels in Western blots. Data are represented as mean ± SEM (n = 3), statistical significance was analyzed by two tail T-test. * represent p<0.05, ** represent p<0.01, *** represent p<0.001, ns represent not significant.

Understanding the biological functions of a given compound without target information, such as a natural product like Vinigrol, is much more difficult and time-consuming, compared to the drug hits originated from the target-driven screening strategy. The functional assessments often require prior knowledge from the bioactivity of their similar compounds. However, this cannot be achieved for a compound with unprecedented backbone structure such as Vinigrol. The fast advance in functional genomics in the past two decades, particularly the transcriptome-profiling technologies, powered with bioinformatic tools, together opened the chance to gain the first unbiased functional prediction of a given compound. This was firstly introduced by the concept named Connectivity Map (CMap), a chemical transcriptome approach brought by Broad Institute in 2006 ^9^. The CMap project developed a database containing large transcriptional expression profiles responding to chemical and genetic perturbagens. The latest version of CMap is called L1000 consisting of >3 million gene expression profiles and >1 million replicate-collapsed gene signatures ^10^, thus creating multiple dimensional connections between genes, chemicals, diseases, and biological status. In the context of pharmacology, CMap is a powerful tool in exploring potential activities of compounds based on the idea that gene expression changes could be used as the universal language to connect the compound without functional annotation to those with known functions. This can be achieved by calculating a “Connectivity Score”, cScore, with an algorithm reflecting the similarities between the inquiry compound to the reference chemicals in CMap database in terms of their gene expression profiles. CMap has been powering both biologists and pharmacists to successfully predict the functions and mechanisms of actions (MOA) of their compounds of interest ^11-13^.

Gene target identification of a compound without functional annotation is another big challenge in pharmaceutical and pharmacological study. The new generation of gene editing technology, CRISPR (Clustered Regularly Interspaced Short Palindromic Repeats), has revolutionized the entire life sciences within only a few years ^14^. Together with CRISPR-associated nuclease (Cas9), which is directed to its target DNA sequence by a short RNA fragment named guide RNA (gRNA), the CRIPSR/Cas9 complex can cut its DNA target from virtually any genome. Consequently, a DNA indel can form at the chosen genomic site, resulting in a frameshift of the mRNA transcript and subsequent nonfunctional protein. This system is referred as CRISPR knockout (CRISPRko). Additionally, scientists engineered a enzymatically inactive (“dead”) versions of Cas9 (dCas9) to eliminate CRISPR’s nuclease activity, while preserving its ability to target desirable sequences. Together with various transcriptional regulators fused to dCas9, it is possible to turn almost any gene on or off or adjust its level of transcription ^15-17^. The dCas9□based transcriptional inhibition and activation systems are commonly referred to as CRISPRi and CRISPRa, respectively. These techniques have a wide range of applications, including for example disease elimination, creation of hardier plants, fight against pathogens and more. In the field of pharmaceutic research, CRISPR has demonstrated an unprecedented power to quantitatively pinpoint both sensitive and resistant gene targets of a drug out of the whole genome gRNA screening and thus unraveled its MOA ^18-20^. When applied with a single gene gRNA, CRISPR emerges as a gold standard to assert a functional gene target of a given compound/drug, through inspecting the activity alterations upon gene manipulation ^21^. CRISPR has brought us into a new era of chemical-genetics for targets-oriented drug discovery.

Endoplasmic reticulum (ER) is a eukaryotic cellular organelle essential for the production, processing, and transport of proteins and lipids. ER dysfunction or stress leads to accretion of misfolded proteins at ER and subsequent initiation of an evolutionarily conserved signaling response, namely unfolded protein response (UPR). Under several physiological and pathological stresses, cells use UPR to sense the pressure and transduce the stress signal through several distinct pathways to downstream transcriptional and post-transcriptional regulations ^22,23^. Consequently, UPR can exert cytoprotective effects by restoring ER function and cellular homeostasis. However, excessive ER stress can also activate intrinsic cellular pathways, leading to cell death ^24,25^. In vertebrates, activation of UPR involves the stimulation of three major signal transducers locating on the ER membrane, i.e., protein kinase R-like ER kinase (PERK), inositol-requiring enzyme 1 alpha (IRE1α), and activating transcription factor 6 (ATF6) (Fig. 2A) ^23,25,26^.

**Figure 2.**
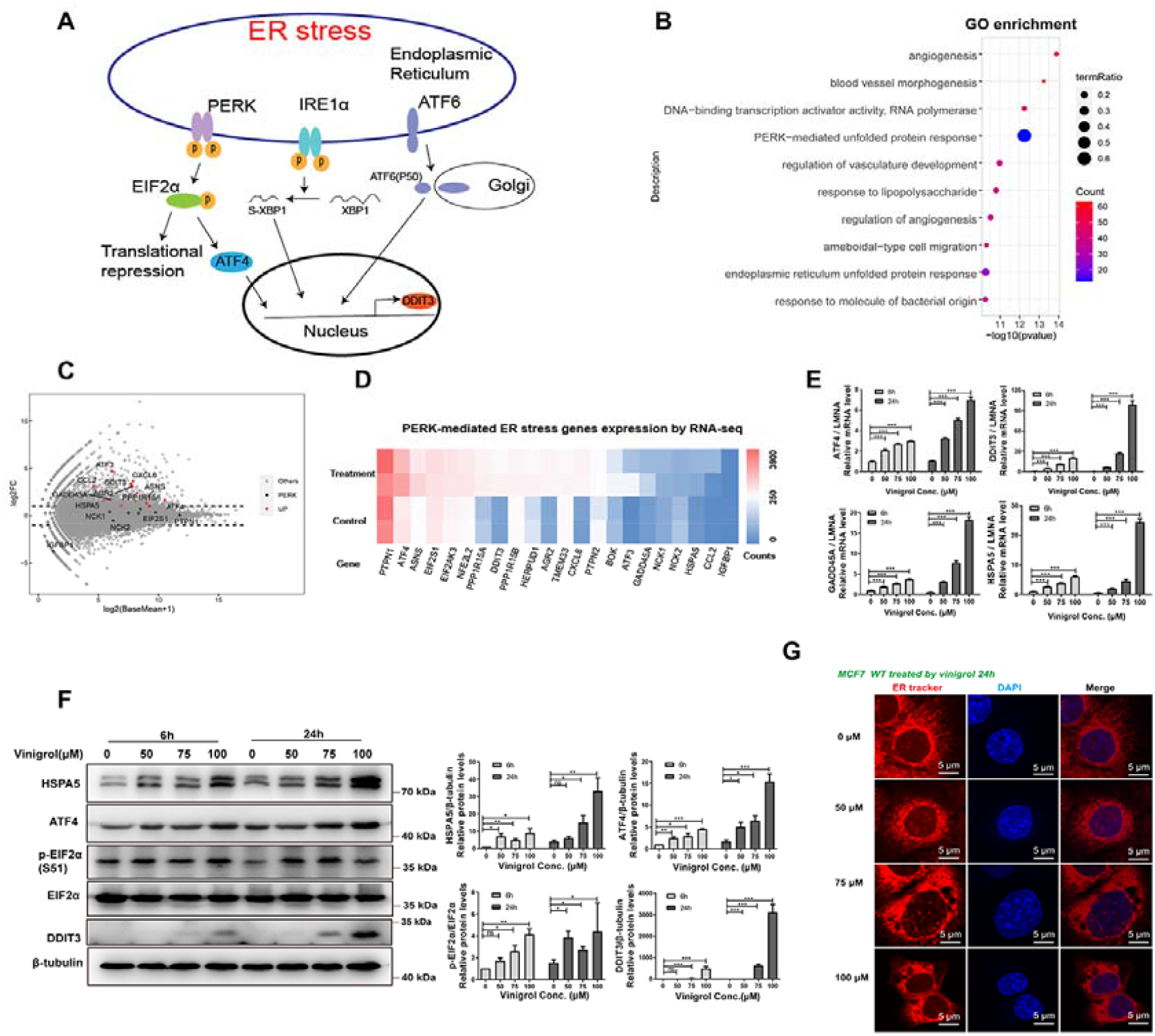
Vinigrol induces PERK mediated ER stress in MCF7. (A) 3 branches of ER stress pathway. (B) Gene Ontology analysis indicates that genes upregulated by Vinigrol are significantly enriched in the term “PERK-mediated unfold protein response”. (C) MA-plot for gene expression comparison between Vinigrol (50μM, 6 hrs) treated for and control MCF7 cells. Red and Black dots denote genes on PERK-mediated ER stress signaling pathway with significant expresson changes (Red) and no changes (Black). (D) Heatmap representing normalized read counts of genes on PERK-mediated ER stress signaling pathway from RNA-seq data. (E) mRNA expression levels of ER stress-related genes with different doses of Vinigrol treatment for different time, quantified by qRT-PCR. (F) (Left) Protein levels of ER stress-related genes with different doses of Vinigrol treatment for different time, quantified by western blot. (Right) Semi-quantification of protein levels. (G) Visualization of ER and nuclei with ER Tracker and DAPI in MCF7 cells after different doses of Vinigrol treatment 24 hrs. Data are represented as mean ± SEM (n = 3), statistical significance was analyzed by T-test. *p<0.05, **p<0.01, ***p<0.001, ns represent not significant.

PERK is a signal-sensitive type I transmembrane protein, which forms a homodimer and trans-autophosphorylate when exposed to ER stress ^27^. The most crucial PERK phosphorylation substrate found so far is eukaryotic translation initiation factor 2 (eIF2α). Upon UPR activation, PERK phosphorylates serine 51 of eIF2α, which subsequently leads to a pause in overall protein synthesis in eukaryotic cells ^27^ but selectively allows translation of activating transcription factor 4 (ATF4), whose target genes include DNA Damage-Inducible Transcript 3 (DDIT3). Under a sustained ER stress response, ATF4 and DDIT3 contribute to the induction of cell apoptosis and autophagy ^28^. On another path, IRE1α, a type 1 ER transmembrane serine/threonine kinase also containing an endoribonuclease (RNase) domain, initiates the most conserved UPR signaling branch ^25^. Under ER stress, IRE1α dimerizes and autophosphorylates to elicit its RNase activity ^29,30^, thereby leading to a catalytic excision of 26-nt intron within the mRNA encoding the transcription factor X-box-binding protein 1 (XBP1) ^31-33^. This stress-stimulated splicing event generates a stable and frameshifted translation product known as XBP1s, the active form of the transcription factor that controls the expression of downstream UPR effector genes ^34^. In the third branch of UPR signaling, full-length ATF6 translocates from the ER to the Golgi apparatus, where it is cleaved by site-1 protease (S1P) and site-2 protease (S2P) to release its C-terminal fragment ATF6f ^35,36^. As an active transcription factor, ATF6 fragment can enter the nucleus and upregulate the expression levels of target genes including specific components in the ER-related protein degradation system and XBP-1^25,31^. In a recent report, both ATF6 and XBP1 are postulated to inhibit colorectal cancer cell proliferation and stemness through activating PERK signaling ^37^. Overall, the three arms of UPR act in parallel and may also combinatorically to maintain ER proteostasis and sustain cell function under ER stress. However, whether all the three brunches are equally important to regulate the stress-induced cell death is far from clear.

In this study, we firstly sought to unbiasedly predict the complete bioactivities of Vinigrol using CMap and confirmed its broad anti-cancer effect on various cancer cell lines. Mechanistically, ER stress turned out as the most affected pathway by Vinigrol. Among the three UPR signaling branches, we demonstrated PERK/eIF2α axis but not IRE1α and ATF6 as the essential pro-death pathway. In this process, ATF4/DDIT3 function as the essential mediators targeted by Vinigrol, as cross-validated by CRISPR/mediated gene repression and activation. Together, our study provided the first mechanistic evidence for the anti-cancer effect of Vinigrol, bringing the whole compound family as novel anti-cancer drug hits.

## 2. Materials and methods

### 2.1 Reagents

Vinigrol (purity 99%) was purified from Fungus was provided by group of Xinsheng Yao and Hao Gao at Jinan University (Guangzhou, China). CellTiter-Glo Luminescent Cell viability (CTG) assay (G7573) was from Promega (Wisconsin, USA). 5x protein loading buffer (P0283), BeyoClick™ EdU-488 assay (C0071), BeyoClick™ EdU-594 assay (C0078), and ER tracker red (C1041) were from Beyotime (Jiangsu, China). Anti-HSPA5(WL03157), anti-ATF4 (WL02330), and Enhanced chemiluminescent plus reagent kit (WLA006C) were from Wanleibio Co., Ltd. (Liaoning, China). Anti-DDIT3 (15204-1-AP) was from Proteintech (Wuhan, China). Anti-β-tubulin (HC101-02) was from Transgenbiotech (Beijing, China). Anti-p21(#2947), anti-CyclinD1 (#2926), anti-CDK2 (#2546), anti-eIF2α (#5324), anti-p-eIF2α (phospho Ser51) (#3597) were from Cell Signaling Technology (Danvers, USA). Anti-p-CDK2 (phospho Thr-14) (ab68265) was from Abcam (Cambridge, UK). Anti-p53(SC-126) was from Santa Cruz (Dallas, USA). Trizol reagent (#R401-01) was from Vazyme (Jiangsu, China). 1640 medium (C11875500BT), FBS (#10270106), and Penicillin-Streptomycin (15140-122) was from Gibco (New York, USA).

### 2.2 Cell culture and cell viability

Human breast cancer cells MCF7 were from ATCC. It was maintained in 1640 medium with 10% FBS in an incubator with 5% CO_2_. Culture media were supplemented with penicillin and streptomycin. The culture medium was changed every two days. MCF7 cells at 70% confluence were used for the next experiment. CTG assay was used to measure cell viability. 10,000 cells per well were seeded in a 96-well plate with 0.1 ml 1640 medium. After 72 hours of drug exposure, 50 μl CTG buffer was added to each well to measure cellular ATP levels. The 96-well plates were shaken 3 min to get cell lysis and then centrifuge at 1000g, 3 min. Luminescent signals were stabilized for 10 min at 25□ and recorded. Each well collection was about 100ms.

### 2.3 RNA sequencing and data analysis

Total RNA was extracted from the MCF7 cells by Trizol reagent. Next-generation sequencing of mRNA-derived cDNA libraries was performed on the Illumina HiSeq X Ten platform at Novogene (Tianijng, China). Control and Vinigrol (50 μM) treatment group were each sequenced with two replicates. At least 9 million reads per sample were generated. All sequencing data were aligned to the GRCh38 genome using the HISAT2 package ^38^. Unique aligned reads were retained to obtain reads count matrices with feature Counts ^39^. The Gene Transfer Format (GTF) file used for quantitative analysis is Ensembl release 90. The Lexogen QuantSeq 3’mRNA-seq library kit (Lexogen, Vienna, Austria) was used to prepare the RNA-seq library in this study. To avoid false increase of gene expression by internal poly-A on the transcripts, we modified the GTF file before generating reads count matrices. The modification is subjected to the last base position of the last exon of each gene in the reference genome. The original sequence of the gene is trimmed to the flanking region of this base. The region of shortened gene body contains 1000nt from this base on five prime directions and 500nt on three prime directions. It is worth noting that the extension on five prime only includes the coding sequence region. In addition, if the length of the original transcript is shorter than 1000nt, the gene is not trimmed. In another direction, if the interval between two genes is smaller than 500nt, the extension on three prime directions is equal to the interval.

Gene expression differences between samples treated by Vinigrol and their corresponding control samples were analyzed by the Bioconductor package DESeq2 ^40^. Fold change value 2.0 and p-value smaller than 0.01 were used as thresholds to distinguish dysregulated genes. Those genes are considered as gene signatures to perform CMap analysis. In this work, the version of the reference database is CMap build02. To input gene signature produced by RNA-seq technique, 22283 Affymetrix IDs in CMap build02 were converted to 13714 gene symbol IDs. A local CMap analysis system was set up to perform analysis. Additionally, both Gene Ontology and KEGG enrichment analysis were performed to deduce the potential biological functions by a Bioconductor package, ClusterProfiler ^41^. Genes with at least one read in treatment or control samples were considered as the enrichment analysis background.

### 2.4 EDU Cell cycle assay

MCF7 cells were treated with Vinigrol or DMSO for 22h, followed by treatment of 10μM 5-ethynyl-2-deoxyuridine (EdU). Three hours after the EdU treatment, cells were harvested and analyzed for EdU signaling using the BeyoClick™ EdU Cell Proliferation Kit with Alexa Fluor 488 according to the manufacturer instructions. EdU reaction buffer (CuSO4, Alexa Fluor 594 azide or 488 azides, and additive buffer) was added to the cells for Click-iT reaction and then incubated for 30□min at 25□ in the dark. Cells were then washed with 3% BSA/DPBS. After click reaction, cells were analyzed on a flow cytometry instrument.

### 2.5 Western blot analysis

Proteins were extracted by RIPA lysis buffer (150mM NaCl, 50mM Tris°HCl, pH 7.4, 1% Triton, 1mM EDTA, 1% sodium deoxycholate, and protease inhibitor mixture) treatment for 15 min in MCF7 cells. Supernatants were collected and centrifuged at 13,000 rpm for 10min at 4□. BCA assays were used to quantify protein contents. Protein samples were then diluted with 5x loading buffer and incubated at 96□ for 10 min in a heating block. Protein samples were separated by 10% SDS-PAGE gel and then electrically transferred to 0.45μm PVDF membranes (Millipore, IPVH00010). Membranes were then blocked with 5% non-fat milk in TBST for one hour at room temperature. Proteins were detected by incubation with indicated primary antibodies overnight at 4°C followed by incubation with HRP-conjugated secondary antibodies for one hour at 25°C. Membranes were subsequently incubated with ECL reagent for 1-2 min and exposed at BIO-RAD ChemiDoc TM XRS+ system (California, USA). All blots were stripped and reblotted with anti-β-tubulin or anti-β-actin as loading controls. All signals were obtained in the linear range for each antibody, quantified using ImageJ pro-plus, and normalized to β-tubulin. The antibodies and the dilutions for western blots used in these studies are as follows: eIF2α (1:2000), p-eIF2α S51 (1:500), HSPA5 (1:2000), and ATF4 (1:1000). DDIT3(1:500), p53(1:300), p21 (1:1000), CDK2(1:1000), p-CDK2 Thr14(1:2000), CyclinD1(1:500), β-tubulin (1:5000).

### 2.6 Quantitative real-time PCR

Total RNA (500 ng) was extracted from the MCF7 cells by Trizol Reagent. PCRs were performed to detect ER stress-related genes. cDNA was synthesized with the HiScript II 1st Strand cDNA Synthesis Kit (Vazyme, R211-01). qPCRs were performed with Hieff qPCR SYBR Green Master Mix (Yeasen, #11201ES08). qRT-PCR conditions were as follows: 95 °C for 5 min, 40 cycles of amplification at 95 °C for 15 s and 60 °C for 1 min, analyzed on the BIO-RAD real-time PCR system. The normalized expression of the assayed genes relative to LMNA was computed for all samples. Primers used to detect the assayed genes are shown in Table S1.

### 2.7 ER tracker staining

Cells were grown overnight on coverslips in 24-well plates. At 24h after treatment with Vinigrol, cells were incubated with ER-Tracker red for 15 min at 37°C. Stained cells were then washed twice with PBS and fixed with 4% formaldehyde for 30 minutes at 25□, followed by hoechst33342 staining and visualized using confocal microscopy (Nikon A1R), using an emission wavelength of 594 nm.

### 2.8 Establishment of ATF4/DDIT3 repression and activation cell lines

To generate the MCF7 cell line stably expressed dCas9-KRAB-MeCP2, approximately 500,000 cells in 6-well plates were transfected with 2000ng dCas9-KRAB-MeCP2-containing PiggyBac expression plasmids (Addgene plasmid #110821) and 400ng of transposase vector PB200PA-1 using PEI. The medium was changed after 24 h. Two days later, cells were treated with 5 μg/ml blasticidin (Yeasen, #60218ES10). Cells were passaged in blasticidin medium for more than two weeks to select dCas9 repressor integrant-containing cells. Single clones were obtained with FACS and the highest inhibition clones were chosen for qRT-PCR verification. We established plasmids contains gRNA ATF4 (F: GCATGGCGTGAGTACCGGGG, R: CCCCGGTACTCACGCCATGC), gRNA DDIT3 (F: GGACCGTCCGAGAGAGGAAT, R: ATTCCTCTCTCGGACGGTCC), and gRNA negative control (NC) (F: GACGACTAGTTAGGCGTGTA, R: TACACGCCTAACTAGTCGTC). After virus packaging, selected single clones were transfected with gRNA vectors and further selected by 1□μg/ml puromycin (MCE, HY-15695). Inhibition efficiency was verified with qRT-PCR.

For CRISPR acitivation (CRISPRa) system, MCF7 cells were transfected with a lentiviral vector expressing dCas9-SunTag10x_v4–P2ABFP-NLS, BFP-positive cells were sorted with FACS. These cells were subsequently transfected with a lentiviral vector expressing scFv-GCN4-GFP-VP64, and GFP-positive cells were sorted with FACS. Single colonies with strong transcriptional activation were sorted for CRISPRa with FACS. Plasmids contain gRNA ATF4 (F: GTAAACGGTTGGGGCGTCAA, R: TTGACGCCCCAACCGTTTAC), gRNA DDIT3 (F: GCCCTAGCGAGAGGGAGCGA, R: TCGCTCCCTCTCGCTAGGGC), and gRNA NC (F: GAACGACTAGTTAGGCGTGTA, R: TACACGCCTAACTAGTCGTTC) were established and their activation efficiency was verified with qRT-PCR.

### 2.9 Internally controlled growth assay

MCF7 CRISPRi cells with stable repression of ATF4, DDIT3, and NC conjugated with RFP reporter and cells with NC gRNA but no RFP-tag were constructed. RFP-positive cells were mixed with RFP-negative cells and were co-cultured for three days. Flow cytometry was used to analyze the proportional change of RFP-positive cells after repeated Vinigrol treatment. In this internally controlled growth assay ^18^, changes in the proportion of RFP-positive cells indicated proliferation speed variation between RFP-positive cells and RFP-negative cells. Proportion of RFP-positive cells was normalized with the proportion of RFP-negative cells compared with the normalized proportion at the original time point. The ratio for each time point was normalized to the same percentage for DMSO control cells, defined as relative enrichment. Therefore, a relative enrichment >1 indicates that RFP positive cells confer protection against treatment, while enrichment <1 indicates sensitization.

### 2.10 Establishment of ATF6 knockout cell lines

The Cas9 stable-expressing MCF7 cells (MCF7-Cas9) were established by lentiviral transfection. Cas9 coding sequence were integrated into the genome of cells (Addgene#52962) and selected with 1 μg/mL blasticidin for nine days. Knockout efficiency was tested using positive control gRNA and indels were confirmed by PCR and DNA sequencing. Knockout cell lines were established by overexpressing gRNA lent guide-Puro (Addgene #52963). After 48h of viral transfection, cells were selected with 1□μg/ml puromycin. Single cells were seeded in individual wells of a 96-well plate and cultured for two weeks. At least 24 individual cell colonies were marked and transferred into a 24-well plate until a sufficient number of cells were obtained to extract genomic DNA. Indels and frameshifts were confirmed by PCR and DNA sequencing. Finally, target gene expression levels were measured with western blot. The ATF6 gRNA sequences are listed as follows: TTAGCCCGGGACTCTTTCAC.

### 2.11 Determination of XBP1 splicing by RT-PCR

The XBP-1 mRNA splicing in MCF7 cells was analyzed by PCR ^42^. The primers’ sequences for PCR amplification of human XBP-1 are 5’-GAATGAAGTGAGGCCAGTGG-3’ and 5’- GGGGCTTGGTATATATGTGG-3’. PCR products were subjected to a 3 % agarose gel with Star Green DNA dye (GenStar, Cat. E107-01) by electrophoresis.

## 3. Results

### 3.1 Vinigrol shows anticancer potency via DNA damage and cell cycle arrest

We firstly applied CMap analysis, a chemical transcriptome-based bioinformatic approach, to explore possible biological activities of Vinigrol. 3’ RNA-seq was performed to obtain the gene expression profiles of Vinigrol-treated MCF7 cells and the differentially expressed genes (DEGs) were identified (see Method). Overall, more than 700 and 330 genes were significantly up- and down-regulated, respectively (Fig. 1B). The DEGs was used as the “gene-signature” to calculate the cScores in comparison to the 6100 transcriptome profiles retrieved from CMap build02 database, which is produced from the cancer cells treated with 1309 compounds/drugs with known biofunctions. In total, 6100 cScores were obtained, normalized and ranked from +1 to -1 (Table S4), reflecting the highest expression similarity (cScore at +1) and dissimilarity (cScore at -1), respectively. Among all the 6100 profiles, those with top positive cScores are enriched for the approved anti-cancer drugs and cytotoxic compounds, suggesting anticancer as the most prominent biofunction of Vinigrol (Fig.1C). Next, we verified the toxicity of Vinigrol on human breast cancer cell line MCF7 (Fig.1D) and 8 other cancer cell lines and identified its broad-spectrum anticancer activity with IC50 at 20-60 μM (ATP measurement with CTG assay) (Table S2). Toxicity of Vinigrol on noncancerous cells was significantly slighter compared to cancer cells (Table S2). Phenotypically, Vinigrol treatment within only 24 hrs could induce severe cytoplasmic vacuolization around nucleus in MCF7 cells at middle to high doses before complete shrinkage and death (Fig. 1E).

We continued the mechanistic studies of Vinigrol on MCF7 cell lines. Firstly, using EdU/hoechst33342 DNA labeling assay followed by FACS flow cytometry quantification, we observed that the proportion of EdU-positive cells, i.e. cells at S phase, were largely reduced whereas the percentage of cells at G2/M phase significantly increased, indicating a cell cycle arrest at G2/M phase (Fig. 1F-1G). This effect of Vinigrol was in a dose-dependent manner, cumulating a complete loss of DNA replication at more than 75 uM of Vinigrol. To dig out the arrest regulators, we further checked the Cyclin/Cyclin-dependent kinase (CDK) cell cycle checkpoints and p53/p21 DNA damage checkpoints (Fig. 1H). Western blot results showed that Vinigrol upregulated p53 and p21, indicating a fast activation of cellular DNA damage response (DDR). Simultaneously, CDK2 was inactivated as demonstrated by a dramatic increase of phosphorylation on its inhibitory site Thr-14. Accordingly, the cell cycle regulator cyclin D1 was severely downregulated. Altogether, it demonstrated that Vinigrol could induce DNA damage and activation of classic p53/p21 DDR pathway, which subsequently inactivated CDK2/Cyclin D1 checkpoints, blocked the replication process, and prevent the cells from moving forward to division. Notably, such effect came out fast and intensified in a time- and dose-dependent manner.

### 3.2 Vinigrol induces PERK-mediated ER stress in MCF7

In light of the Vinigrol’s anticancer effect, we performed gene enrichment analysis to explore its potential molecular mechanisms. The up- and down-regulated genes upon Vinigrol treatment were subjected to gene ontology (GO) analysis, respectively, using all expressed genes as background (Fig. 2B). GO enrichment analysis suggested that many of the upregulated genes are involved in PERK-mediated ER stress (Fig. 2B-2D). Strong and extended ER stress can result in cell dysfunction and even cell death ^24,25^. Quantitative Reverse Transcription PCR (qRT-PCR) measurements revealed continuous rises of mRNA level of the key PERK-mediated ER stress genes, including stress initiator HSPA5/Bip, intermediator ATF4, and downstream death effector DDIT3 and GADD45A (growth arrest and DNA damage inducible 45 alpha), along with increase of drug doses and treatment duration (Fig. 2E). Consistent with the mRNA changing dynamics, protein levels of hallmark molecules including HSPA5/Bip, p-eIF2α, eIF2α, ATF4, and DDIT3, also sustainably increased (Fig. 2F, Fig.S1). These data showed that Vinigrol can induce a fast activation of ER-stress sensor HSPA5 and PERK arm of UPR, which subsequently phosphorylates eIF2α, leading to induction of downstream transcriptional regulator ATF4 and pro-death effector DDIT3. Notably, the ER stress activation by Vinigrol was in a time- and dose-dependent manner, whose dynamics was in accordance with the changes of p53/p21 DNA damage responders and cell cycle checkpoints (Fig. 1G), suggesting additive role of ER stress in regulating the cell death. This was further confirmed by reduced cell valibity through combined treatment of PERK-specific inhibitor GSK2606414 with Vinigrol (Fig. S2). When we labeled the ER of MCF7 cells with fluorescent dye ER Tracker Red, we observed that the ER gradually lost its structure, enlarged and vacuolized after Vinigrol treatment. Vacuolization in the ER was also exacerbated with higher doses of Vinigrol (Fig. 2G). ER dilation and cytoplasm vacuolization are key morphological features of ER stress. Altogether, these results showed that Vinigrol could induce cell deaths by activating PERK-mediated ER stress pathway.

### 3.3 Vinigrol induced ER stress can be alleviated by DDIT3 and ATF4 knockdown

PERK-mediated ER stress is related to increased abundance of ATF4, a transcriptional factor that directly activates DDIT3 transcription ^28^. To determine whether Vinigrol induces cell death is via the ATF4-DDIT3 axis, we developed cell lines with stable repression of ATF4 and DDIT3 using CRISPR/dCas9 inhibition (CRISPRi) system ^16^. The knockdown efficiency of target genes was verified by qRT-PCR (Fig. 3A), which demonstrated that mRNA expression levels of ATF4 and DDIT3 were significantly repressed in MCF7 cells comparing with the dummy guide RNA (gRNA)-infected negative control (NC) MCF7 cells. Since ATF4 and DDIT3 are key intermediate regulator on PERK arm of UPR signaling, we also checked the mRNA and protein expression of other upstream and downstream regulators HSPA5/BiP, eIF2α and GADD45A as did on MCF7 wildtype. Upon Vinigrol treatment, repression of either ATF4 or DDIT3 could significantly alleviate the drug-induced upregulation scale of HSPA5/Bip and GADD445A mRNA level, albeit not fully rescued the increase (Fig. 3B). Such alleviation was also observed on the protein level. Comparing to NC-MCF7 wildtype cells, the upregulation of both HSPA5/Bip and p-eIF2α were almost completely abolished in ATF4 or DDIT3 CRISPRi repression cells (Fig. 3C). These results indicate an important feedback loop of ATF4 and DDIT3 to retro-regulate their upstream UPR mediators such as HSPA5/Bip and eIF2α, beyond transducing stress signals to their downstream targets. Interestingly, repression of ATF4 largely refrained DDIT3 from being stimulated by Vinigrol-induced ER stress at both mRNA and protein level (Fig. 3B-C). On the other hand, DDIT3 repression barely influenced the ATF4 transcription and translation (Fig. 3B-C), even though the gRNA targeting on DDIT3 provided higher knockdown efficiency than ATF4 gRNA (Fig. 3A-B). These data further confirmed DDIT3 as the transcriptional target of ATF4, but not vice versa. Consequently, ATF4 functions on top of DDIT3 to promote the PERK signaling. Both transcriptional factors, however, play essential roles in retaining the Vinigrol-induced ER stress responses.

**Figure 3.**
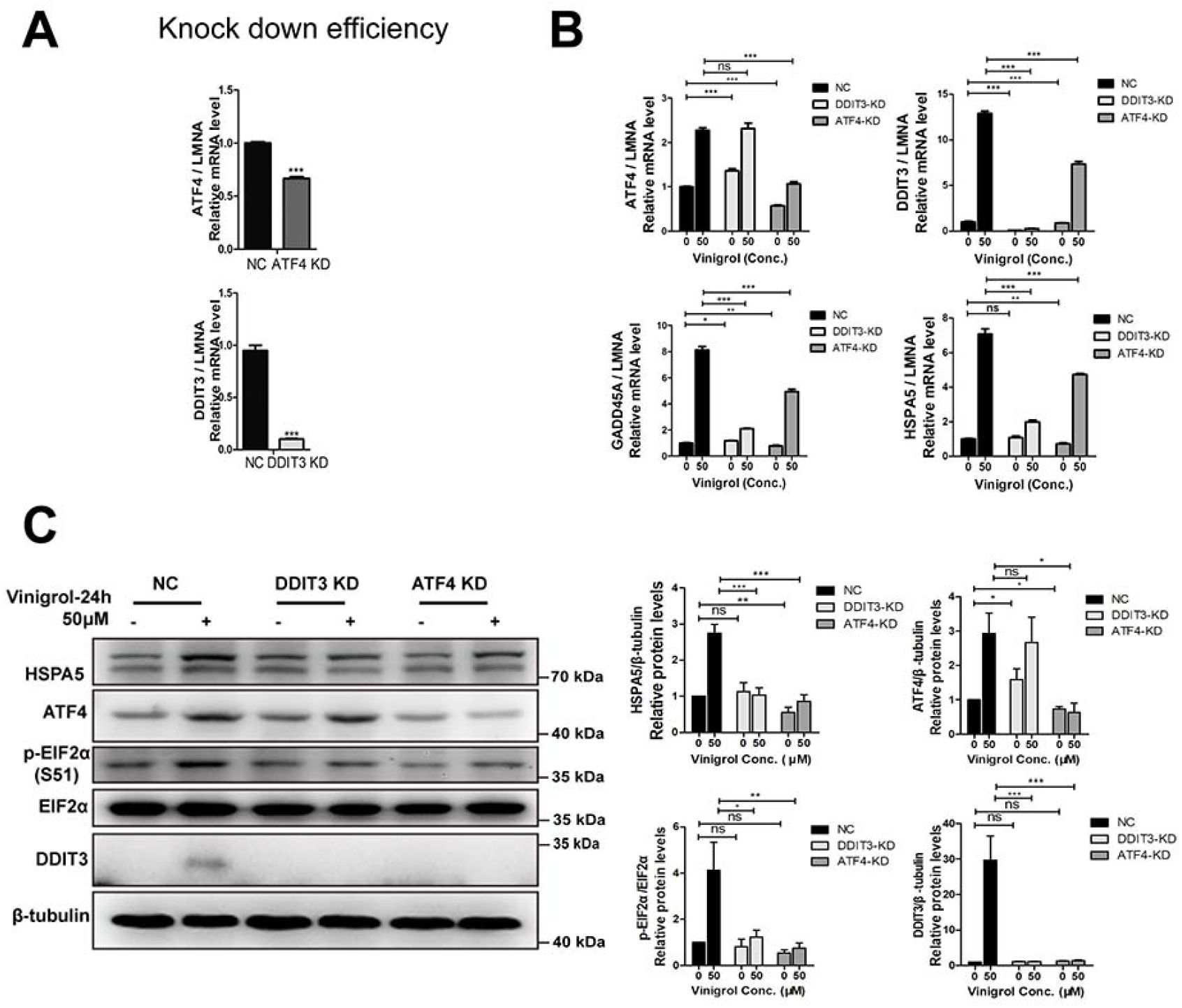
ER stress-induced by Vinigrol was alleviated after ATF4/DDIT3 knockdown. (A) Knockdown efficiency of ATF4 and DDIT3 in MCF7 by CRISPR/dcas9 inhibition system, *compared with NC, ***p<0.001. (B) mRNA expression levels of ER stress-related genes after 24 hrs of Vinigrol (50μM) treatment, quantified by qRT-PCR. (C) (Left) Protein levels of ER stress-related protein after Vinigrol (50μM) treatment for 24 hrs, detected by Western blot. (Right) Semi-quantification of protein levels. Data are represented as mean ± SEM (n = 3), statistical significance (p<0.05) was analyzed by T-test. *p<0.05, **p<0.01, ***p<0.001, ns represent not significant.

### 3.4 Vinigrol-induced cell death can be alleviated by ATF4/DDIT3 knockdown

Stress induced UPR can contribute to cellular homeostasis and recovery, and cell death. We therefore investigated the concrete role of ATF4/DDIT3 in Vinigrol-induced death process. Cell viability measurement (CTG assay) revealed a clear rescue of ATF4 or DDIT3 knockdown on cell growth when treated with Vinigrol at the concentrations near IC50 (Fig. 4A). The growth protection by knocking down ATF4 and DDIT3 were further supported by EDU/hoechst33342 cell cycle analysis. Comparing to the NC cells with dummy gRNA, repression of the genes retained the DNA replication of the cells, as demonstrated by increased percentage of S-phase cells (Fig. 4B-C). Notably, such increase was only observed upon Vinigrol treatment, not in vehicle treatment controls, which excluded the gene knockdown, by itself, from protecting the DNA replication. Accordingly, when we checked the protein level of p21, a key cell cycle negative regulator and DNA damage checkpoint, repression of ATF4 and DDIT3 specifically diminished the drug-induced upregulation of p21 (Fig. 4D), and thus promoted the cell cycle process.

**Figure 4.**
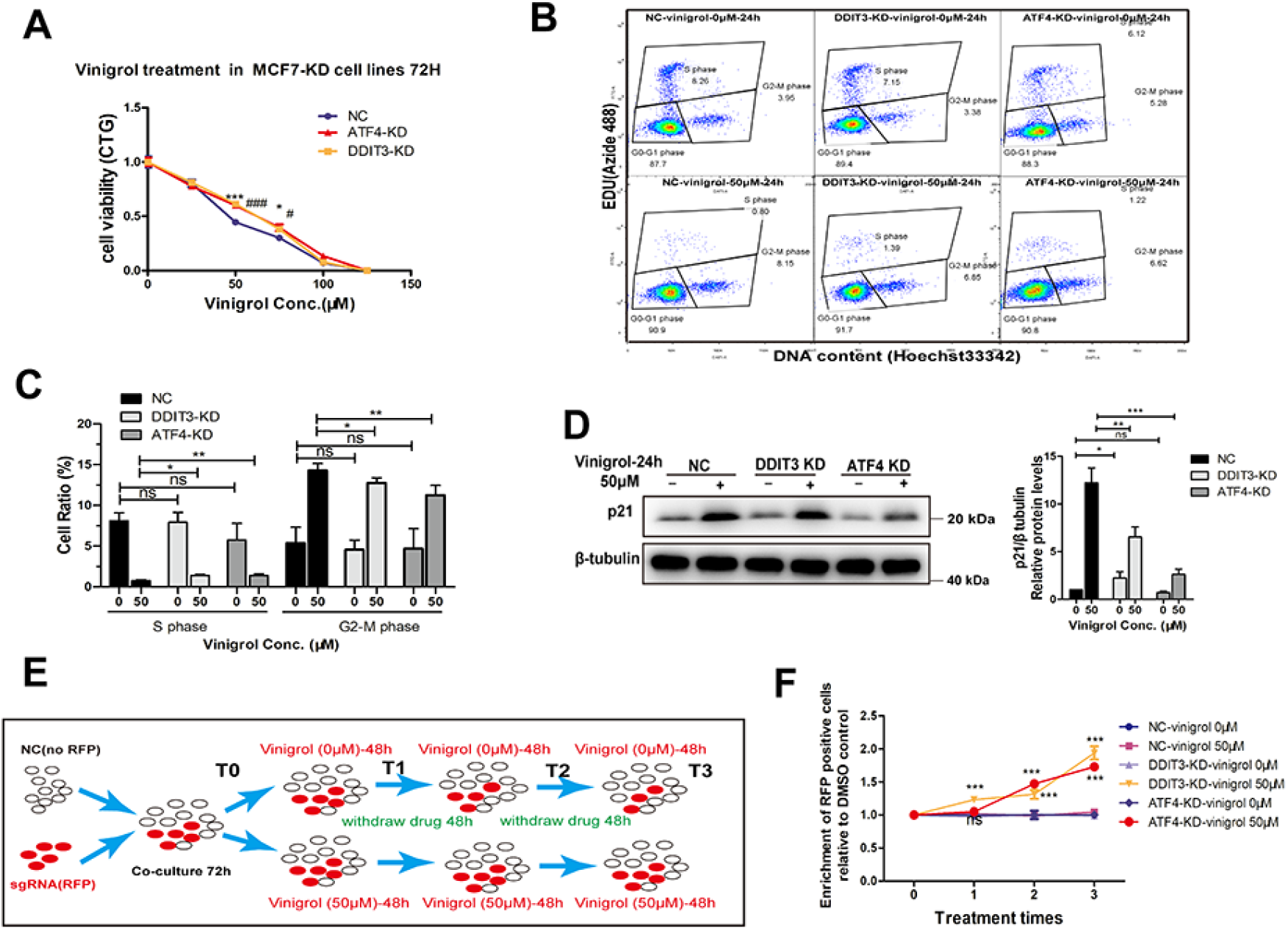
Cell death induced by Vinigrol was alleviated after ATF4/DDIT3 knockdown. (A) Cell viability quantified by CTG assay after Vinigrol treatment for 72 hrs. ATF4-KD group compared with the NC group, # p<0.05, ## p<0.01, ### p<0.001; DDIT3-KD group compared with the NC group, * p<0.05, ** p<0.01, *** p<0.001; ns: not significant. (B) EDU/Hoechst33342 cell cycle analysis after DDIT3 and ATF4 knockdown, analyzed by BD FACS. (C) Quantification of cells in different stages of cell cycle. (D) Protein level of p21 after 50μM Vinigrol treatment for 24 hrs, detected by western blot. (E) Workflow of internally controlled growth assay. (F) Enrichment of RFP positive cells relative to DMSO-treated control cells across three repeated 50μM Vinigrol treatments, analyzed by flow cytometry. Data are represented as mean ± SEM (n = 3), statistical significance was analyzed by T-test, * p<0.05, ** p<0.01, *** p<0.001, ns represent not significant.

To further investigate the effects of ATF4 and DDIT3 knockdown on Vinigrol sensitivity in an extended time frame, we applied an internally controlled growth assay (see 2.9 in Materials and methods). Briefly, in this experiment, the red fluorescent protein (RFP) was introduced to label the ATF4 or DDIT3 gRNA infected cells, which were co-cultured, respectively, with the non-fluorescent dummy gRNA cells (NC) followed by three runs of Vinigrol treatment and withdraw (Fig. 4E). The proportion of RFP to NC cells was quantified after each run of drug treatment by flow cytometry to indicate the relative growth rates of CRISPRi gene edited cells compared to non-edited cells. Such RFP/NC proportion was further normalized by the same co-culture setup but with only DMSO control treatment, in order to exclude the gene effect on the cell growth that is irrelevant to drug effects. Overall, RFP positive cells displayed relative enrichment greater than 1 and keep stepping up with repeated drug treatment, which indicated that repression of ATF4 or DDIT3 confers growth protection against Vinigrol treatment (Fig. 4F). Together, these findings clearly indicate ATF4 and DDIT3 as two essential pro-death mediators that are induced by Vinigrol and directly contribute to its anti-cancer effect.

### 3.5 Vinigrol induced cell death can be exacerbated by ATF4/DDIT3 overexpression

After establishing the necessity of ATF4/DDIT3 axis in Vinigrol induced cell deaths through CRISPRi gene editing system, we wondered if activation of this pathway would exacerbate Vinigrol induced cell death. To this end, we constructed MCF7 cells stably overexpressing ATF4 or DDIT3 transcripts using CRISPR/dCas9-SunTag-VP64 gene engineering system, namely CRISPR activation (CRISPRa) system ^17,43^ coupled with the gRNAs specifically targeting ATF4 and DDIT3, respectively. The overexpression of both genes was efficient as quantified by using qRT-PCR (Fig. 5A). In contrast to the knockdown results, ATF4 or DDIT3 overexpression significantly exacerbated the Vinigrol-induced ER stress as demonstrated by the enlarged upregulation magnitudes of HSPA5/Bip, GADD45A, and p-eIF2a at mRNA level (Fig. 5B) and protein level (Fig. 5C). As expected, activation of ATF4 promoted DDIT3 transcription and led to protein level increase in both Vinigrol and control treatments (Fig. 5B-C). Consequently, CTG cell viability measurement and EdU cell cycle analysis showed that cell death induced by Vinigrol was significantly aggravated after ATF4 or DDIT3 overexpression (Fig. 5D-5F). Specifically, the fraction of S-phase cells was further reduced when overexpressing ATF4 or DDIT3 upon Vinigrol treatment. Notably, without the drug treatment, activation of either gene could significantly promote the cells into S-phase (Fig. 5E-G), indicating strong protective effects on the cell growth. When such basal gene effects were normalized out to obtain the Vinigrol-induced gene effects (by comparing the reduction fold between Vinigrol/DMSO groups), we found that the DNA replication stress induced by Vinigrol was deteriorated with ATF4 or DDIT3 overexpression (Fig. 5F). This was further supported by the enhanced upregulation scale of p21 in CRISPRa engineered cells (Fig. 5G). In the internally controlled growth assay (Fig. 4E), in contrast to the growth rescue from ATF4 and DDIT3 knockdown, ATF4 and DDIT3 overexpression cells were almost depleted after repeated Vinigrol treatment (Fig. 5H). Altogether, our results from both CRIPSR inhibition and activation systems provide solid evidence to pinpoint ATF4/DDIT3 as the functional targets of Vinigrol, whose expression levels dictate the cell sensitivity and resistance to the anti-cancer effect of Vinigrol.

**Figure 5.**
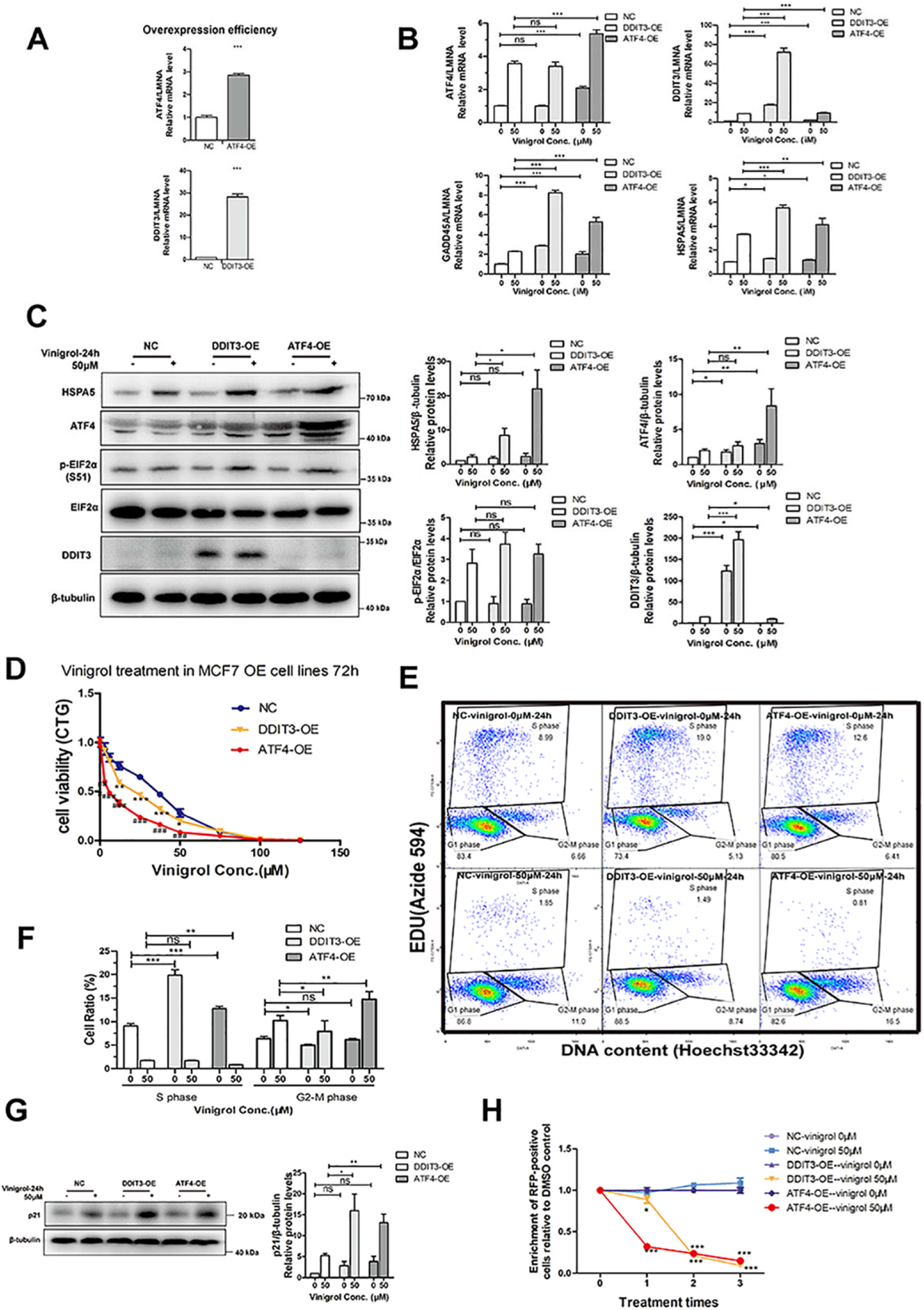
Cell death induced by Vinigrol was exacerbated by overexpression of ATF4/DDIT3. Cell death induced by Vinigrol was exacerbated by overexpression of ATF4/DDIT3. (A) Overexpression (OE) efficiency of ATF4 and DDIT3 by CRISPR/dcas9 activation system, *compared to NC, ***p<0.001. (B) mRNA expression levels of ER stress-related genes after 50 μM Vinigrol treatment for 24 hrs, quantified by qRT-PCR. (C) (Left) Protein levels of ER stress-related genes after 50 μM Vinigrol treatment for 24 hrs, detected by western blot. (Right) Semi-quantification of protein levels. (D) Cell viability after Vinigrol treatment for 72 hrs, quantified by CTG assay. ATF4-OE group compared with NC group, # p<0.05, ## p<0.01, ### p<0.001; DDIT3-OE group compared with NC group, * p<0.05, ** p<0.01, *** p<0.001; ns: not significant. (E) EDU/Hoechst33342 cell cycle analysis after ATF4 and DDIT3 overexpression, analyzed by BD FACS. (F) Quantification of cells in different cell cycle stages. (G) Protein level of p21 after 50 μM Vinigrol treatment 24 hrs, detected by western blot. (H) Enrichment of RFP positive MCF7 cells relative to DMSO-treated control cells in internally controlled growth assay. Data are represented as mean ± SEM (n = 3), statistical significance was analyzed by T-test, * p<0.05, ** p<0.01, *** p<0.001, ns represent not significant.

### 3.6 PERK, but not ATF6 or IRE1α, directly contribute to Vinigrol induced cell death

Among the three ER stress pathways, we next inspected the role of ATF6 and IRE1α pathways in Vinigrol induced cell death. From the RNA-seq analysis, most of ER stress related genes were significantly upregulated upon Vinigrol treatment (Fig. 6A-6B). However, ATF6 and IRE1α pathways were not enriched by GO and KEGG analysis. ATF6 is an essential messenger of ER stress signaling pathway. Previous studies indicated that it is involved in sustaining cell viability ^37^. ATF6 gene expression level increased in a dose- and time-dependent manner after Vinigrol treatment (Fig. 6C). Consistently, ATF6 protein level significantly increased at 6 hrs and quickly decreased at 24 hrs (Fig. 6D). Furthermore, we used CRISPR/Cas9 to generate single clones of MCF7 cells carrying homozygous deletion of ATF6 and treated them with Vinigrol. Unexpectedly, we found that loss of ATF6 did not affect Vinigrol sensitivity compared with controls in CTG cell viability assay (Fig. 6E-F), which suggested that ATF6 is dispensable for Vinigrol-triggered cell death, and the fast increase of ATF6 expression level is likely a temporary response of cells to Vinigrol treatment.

**Figure 6.**
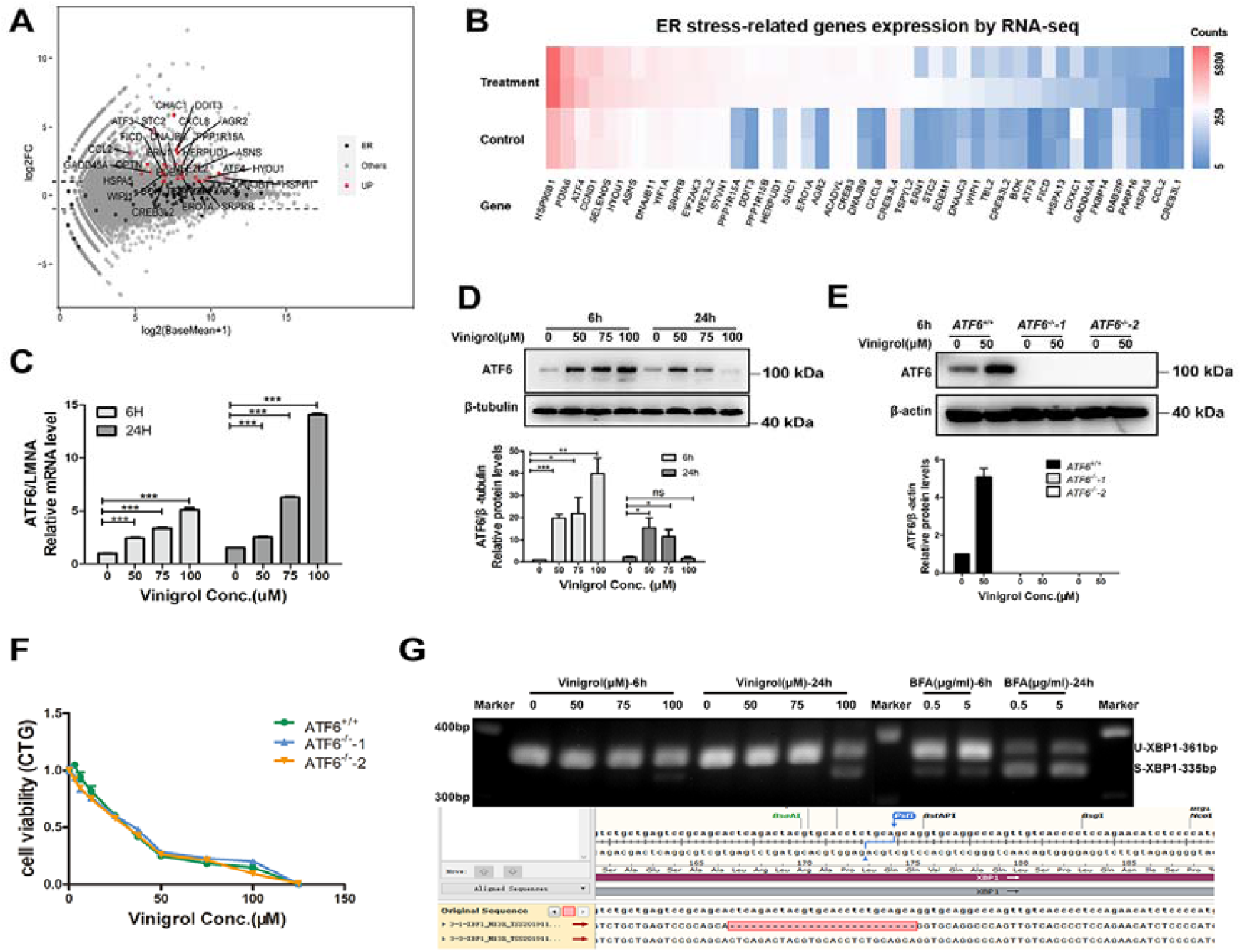
Effect of Vinigrol on ATF6 and IRE1α pathways in MCF7. (A) Expression levels of ER stress-related genes shown in MA plot in RNA-seq analysis. Red and Black dots denote genes on ER stress signaling pathway with significant expression changes (Red) and no changes (Black). (B) Expression levels (normalized RNA-seq read counts) of ER stress-related genes shown in heatmap. (C) mRNA level of ATF6 after varied doses of Vinigrol treatment for different time, quantified by qRT-PCR. (D) Protein levels of ATF6 expression level after Vinigrol treatment in MCF7 cell, detected by western blot. (E) Protein levels of ATF6 after 50 μM Vinigrol treatment for 6 hrs in ATF6^-/-^ MCF7 cell lines, quantified by western blot. (F) Cell viability after Vinigrol treatment in ATF6^-/-^ MCF7 cell lines, quantified by CTG assay. (G) Splicing of XBP1 by IRE1 quantified by RT-PCR, with BFA as a positive control. Sanger sequencing results of the two PCR amplicons revealed 26-nt deletion of s-XBP1. Data are represented as mean ± SEM (n = 3), statistical significance was analyzed by T-test. * p<0.05, ** p<0.01, *** p<0.001, ns represent not significant.

IRE1α/XBP1 pathway plays important roles in UPR. Under extended stress, IRE1α activation can cause cell death by activating the MAPK pathway ^44,45^. Vinigrol treatment did not induce XBP-1 splicing; only slightly spliced-XBP-1 was detected at 100 μM Vinigrol treatment for 24 hrs. Brefeldin A (BFA), a ER stress inducer was known to activate IRE1α to splice XBP-1 ^46^. As a positive control in our study, BFA reapidly led to a fast splicing of XBP-1 at low concentrations in short time (Fig. 6G), which in turn suggested that the IRE1α pathway was not activated in Vinigrol induced cell deaths. To summarize, our data indicated that Vinigrol functions through PERK-mediated ER stress instead of ATF6 and IRE1α pathways to cause cell death in MCF7 cells.

## 4. Discussion

The fundamental goal of this study is to decipher the bioactivity and functional mechanism of Vinigrol, a natural diterpenoid with unprecedented chemical structure. The success of its full synthesis achieved in the past decade not only create a series of compound analogues, but also shed lights on their potential medical use through elaborated pharmacological research and drug development. In the context of molecular biology, we are also intrigued by a more fundamental question: how the structure uniqueness of Vinigrol interferes with the cellular environment of a mammalian cell.

In light of the inventions in transcriptome analytic techniques, i.e., Microarray as the 1st generation of invention and then RNA-seq afterwards, we applied CMap for the first functional assessment of Vinigrol. Up to now, three versions of CMap database have been updated with the number of total expression profiles increasing from a hundred (Build01), 6100 (Build02), and above a million (L1000). Notably, only the first two versions provide the experimentally quantified whole transcriptome datasets that are performed on Affymatrix Microarray. In L1000, on the other hand, the expression profiles of 12291 genes were computationally imputed from the actual quantification of only 978 genes, which would reduce the calculation accuracy of cScore ^10^. To achieve more precise functional predictions, we decided to compare our RNA-seq data of Vinigrol to Build02, whose 1309 perturbagens consist of most of approved drugs and drug hits with annotated actions and molecular targets.

Among the 20 best-matched drugs (Table S3), apart from anti-cancer effect, our results unraveled several other potential biofunctions of Vinigrol, such as anti-hypertension (similar to, for example, adrenergic receptor antagonist drug phenoxybenzamine and suloctidil), anti-depression and antipsychosis (similar to protriptyline, thioridazine, prochlorperazine, perphenazine). Interestingly, the anti-hypertensive activity of Vinigrol was reported along with its discovery in 1988, the only pharmacological study of Vinigrol ever published till now ^7^. In this early study, Vinigrol was orally given to conscious spontaneously hypertensive rats and its anti-hypertensive activity was observed, yet the underlying mechanism and molecular targets were completely missing. Experimentally approved anti-hypertension and anti-cancer (this study) effect of Vinigrol demonstrate the unbiasedness and reliability in functional prediction using chemical transcriptome-based approach. Undoubtedly, the predicted effects of Vinigrol on neuronal and psychotic diseases such as depression, AD, and schizophrenia are in high medical needs and definitely worth of further exploration. However, to find the correct targets perturbation and suitable disease models persists as the leading challenge to scientists, and our CMap results and RNA-seq oriented gene and pathway analyses provide valuable multi-dimensional information to guide their experimental design.

In our gene set enrichment analyses, ER stress/UPR relevant terms stood out as the pathways that are mostly likely targeted by Vinigrol (Fig. 6A). However, all the three UPR branches, PERK/eIF2α, ATF6, and IRE1α are firstly known to be essential in restoring the cellular homeostasis and, when under excessive and prolonged activation, also result in stress-induced inflammation and apoptosis ^24,25,27,28,45,46^. To fully understand the role of ER stress in the anti-cancer effect of Vinigrol, it is necessary to address the question whether the three pathways equally attributes to the stress adaptation of the cell or mediates the cell death in response to Vinigrol. We applied multiple cellular, biochemical, and molecular assays, together with CRISPR/Cas9 mediated gene knockout, repression, and activation systems, to specifically investigate the key mediators regulating each branch. The results clearly showed that Vinigrol-induced cell death largely attributes to the activation of PERK/eIF2α arm instead of IRE1α/XBP1 and ATF6 branches. Despite the IRE1α transcript slightly increases, it fails to cleave the XBP1 RNA and keep the pathway silent. It was reported that proteasome inhibitors can induce ER stress but prevent IRE1α from splicing XBP-1 ^47^. In our CMap analysis, the expression profiles of Vinigrol treatment matched those of celastrol (Table S3), a proteasome inhibitor that prevents XBP-1 splicing. Vinigrol’s proteasome inhibition activity may be the cause of inactivated XBP-1 during Vinigrol treatment. The activation of ATF6 expression was dramatic shortly after Vinigrol treatment, suggesting it also plays a role. However, not until we achieved its homozygous deletion by using CRISPR/Cas9 knockout system could we affirm its irrelevance to the cell growth (Fig. 6F). Thus, we postulate the activation of ATF6 as a downstream stress response rather than a functional mediator of Vinigrol’s anti-cancer effect.

PERK/eIF2α is probably the most profoundly studied branch among the three, involving the translational control by eIF2α that is phosphorylated by PERK, and downstream overexpression of two transcriptional factor ATF4 and DDIT3. Therefore, we carefully inspected the changing dynamics of ATF4 and DDIT3 together with their upstream mediator HSPA5/Bip, eIF2α and several downstream effectors, and revealed that they are all gradually upregulated by Vinigrol in a dose- and time-dependent manner. Taking advantages of CRISPRi and CRISPRa, we cross-validated the essential regulatory roles of ATF4/DDIT3 in promoting the death process stimulated by Vinigrol. Specifically, knockdown of each gene restrains all the studied genes from being over-activated by Vinigrol and thus alleviated the death, whereas CRISPRa exerts the opposite effects. It is worth mentioning that the influential scales of gene overexpression and knockdown sometimes are not at the same magnitude in terms of their functional consequences, even if they are opposite. This is largely depending on the basal expression of a gene. Specifically, in the MCF7 cells growing in normal condition, the basal expression of DDIT3 is barely detected, therefore its further knockdown renders relatively minor influence compared to it overexpression on the cell growth (Fig. 4A, 5D). Thus, it is always encouraged to apply both repression and activation in gene-editing experiments to cross-validate the results. Interestingly, even without Vinigrol treatment, the CRISPRa enhanced basal expression of DDIT3 elicits dramatic increase of S-phase cells (Fig. 5E-F). Previous reports described more of pro-death effect of DDIT3 under ER stress through inducing the expression of pro-apoptosis gene targets such as death receptor 5 (DR5) and Growth arrest and DNA damage-inducible protein 34 (GADD34) ^25,48^. Here, we clearly demonstrated a favorable effect of slight upregulation of DDIT3 to promote the DNA replication of MCF7 tumor cells. However, its excessive and prolonged activation stimulated by chemicals such as Vinigrol, halts the cell cycle and facilitates the death process that is mediated by the best understood regulatory axis p53/p21. Altogether, our study unveiled a broad anti-cancer function of Vinigrol with good potential as a clinical medication. Nevertheless, we strongly recommend monitoring the expression of key regulators such as ATF4 and DDIT3 before and during chemotherapeutic process to guarantee a sufficient activation.

## 5. Conclusion

Vinigrol, a natural diterpenoid with unprecedented chemical structure, has gained a huge endeavor in the past decade for its chemical synthesis. The success of chemists on its full synthesis created a series of compound analogs with similar structures that await functional and mechanistic investigation.

In this study, we firstly applied CMap to unbiasedly predict the complete bioactivities of Vinigrol. Anti-cancer appear to be the most prominent biofunction followed by anti-hypertension and anti-depression/psychosis. A broad cell toxicity on various cancer cell lines was further confirmed. Mechanistically, ER stress turns out as the most affected pathway by Vinigrol. Among the three UPR braches, we demonstrate PERK/eIF2α axis but not IRE1α and ATF6 as the essential pro-death pathway, which subsequently activates the cell cycle negative regulator and DNA damage checkpoints p51/p21. In this process, ATF4/DDIT3 function as the essential mediators targeted by Vinigrol, as cross-validated by CRISPRi/a genetic engineering. In conclusion, we provided the first evidence for the anti-cancer effect of Vinigrol and the underlying mechanisms. With the recent achievement of Vinigrol’s chemical synthesis, our results also paved the way for further comparisons and elucidation of the structure-effect relationship of the whole compound family, making them a novel chemical cluster with potential for cancer therapy. Additionally, this study serves as a valuable resource on the genes and pathways affected by Vinigrol, as well as a complete list of potential bioactivities to facilitate other pharmacological validations.

## Acknowledgments

This work was supported by the Shenzhen Key Laboratory of Gene Regulation and Systems Biology (Grant No. ZDSYS20200811144002008, China) (Yuhui Hu), the Shenzhen Science and Technology Program (Grant No. KQTD20180411143432337, China) (Yuhui Hu), the National Natural Science Foundation of China (Grant No. 81773881, China) (Yuhui Hu). We thank Wei Chen and Qinan Hu (Southern University of Science and Technology, SUSTech, Shenzhen, China) for inspiring discussions, suggestions and comments for the experiment and manuscript preparation. Computational resource and experimental facilities were supported by the Center for Computational Science and Engineering and the Core Research Facilities at SUSTech.

## Supporting information

**Figure S1.**
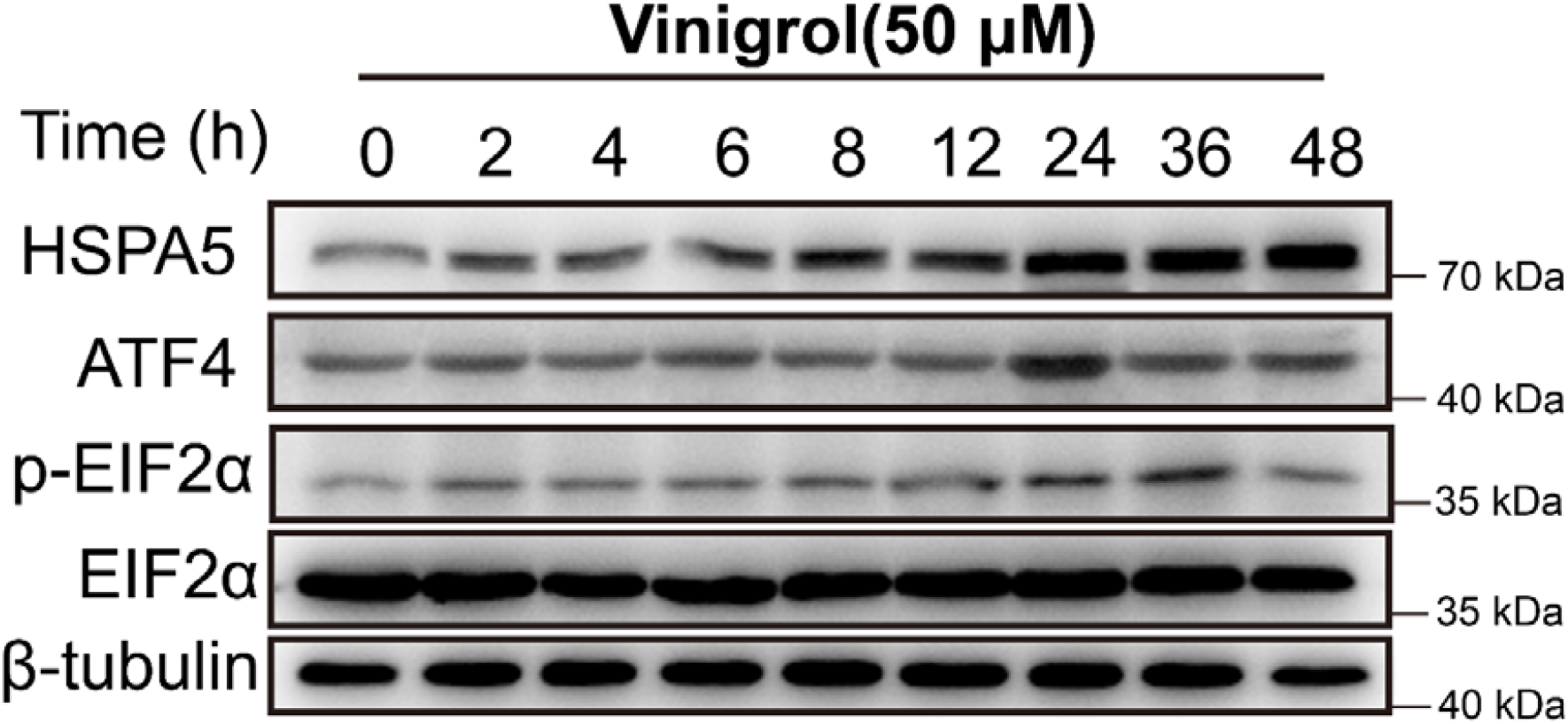
The ER stress-related protein expression level of MCF7 cells after Vinigrol (50μM) treatment at different time.

**Figure S2.**
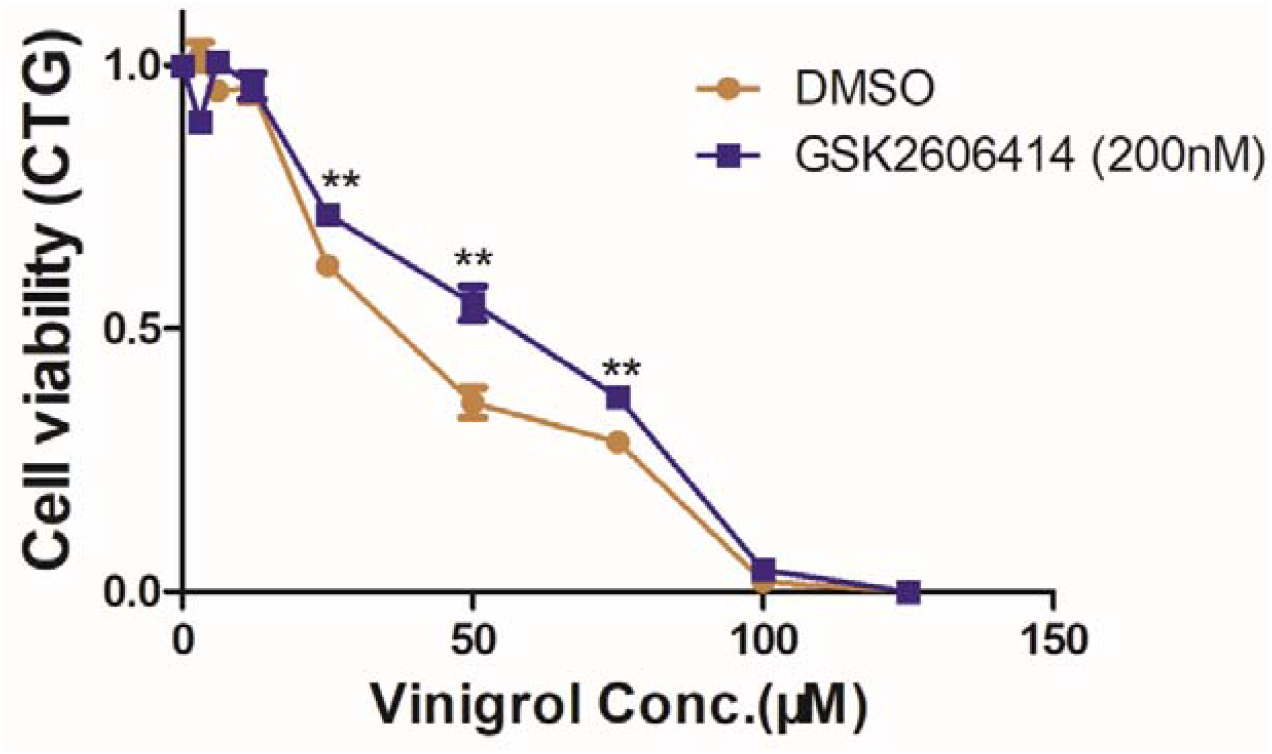
Cell viability of MCF7 cells treated with Vinigrol with and without GSK2606414 (200nM) for 72 hrs.

## References

1 Uchida, I. et al. The structure of vinigrol, a novel diterpenoid with antihypertensive and platelet aggregation-inhibitory activities. The Journal of Organic Chemistry 1987; 52: 5292–5293.

2 Min, L., Lin, X. & Li, C. C. Asymmetric Total Synthesis of (-)-Vinigrol. J Am Chem Soc 2019; 141: 15773–15778.

3 Yu, X., Xiao, L., Wang, Z. & Luo, T. Scalable Total Synthesis of (-)-Vinigrol. J Am Chem Soc 2019; 141: 3440–3443.

4 Gentric, L., Hanna, I. & Ricard, L. Synthesis of the complete carbocyclic skeleton of vinigrol. Org Lett 2003; 5: 1139–1142.

5 Huters, A. D. & Garg, N. K. Synthetic studies inspired by vinigrol. Chemistry 2010; 16: 8586–8595.

6 Poulin, J., Grise-Bard, C. M. & Barriault, L. A formal synthesis of vinigrol. Angew Chem Int Ed Engl 2012; 51: 2111–2114.

7 Ando, T., Yoshida, K. & Okuhara, M. Vinigrol, a novel antihypertensive and platelet aggregation inhibitory agent produced by a fungus, Virgaria nigra. II. Pharmacological characteristics. J Antibiot (Tokyo) 1988; 41: 31–35.

8 Paquette, L. A., Guevel, R., Sakamoto, S., Kim, I. H. & Crawford, J. Convergent enantioselective synthesis of vinigrol, an architecturally novel diterpenoid with potent platelet aggregation inhibitory and antihypertensive properties. 1. Application of anionic sigmatropy to construction of the octalin substructure. J Org Chem 2003; 68: 6096–6107.

9 Lamb, J. et al. The Connectivity Map: using gene-expression signatures to connect small molecules, genes, and disease. Science 2006; 313: 1929–1935.

10 Insititute, B. Connectivity Map, 2006. https://clue.io/.

11 Li, Z. & Yang, L. Underlying Mechanisms and Candidate Drugs for COVID-19 Based on the Connectivity Map Database. Front Genet 2020; 11: 558557.

12 Vanderstocken, G. et al. Identification of Drug Candidates to Suppress Cigarette Smoke-induced Inflammation via Connectivity Map Analyses. Am J Respir Cell Mol Biol 2018; 58: 727–735.

13 Yoo, M. et al. Exploring the molecular mechanisms of Traditional Chinese Medicine components using gene expression signatures and connectivity map. Comput Methods Programs Biomed 2019; 174: 33–40.

14 Jinek, M. et al. A programmable dual-RNA-guided DNA endonuclease in adaptive bacterial immunity. Science 2012; 337: 816–821.

15 Shalem, O., Sanjana, N. E. & Zhang, F. High-throughput functional genomics using CRISPR-Cas9. Nat Rev Genet 2015; 16: 299–311.

16 Joel R. McDade, N. C. W., Lianna E. Swanson, Melina Fan. Practical Considerations for Using Pooled Lentiviral CRISPR Libraries. Current Protocols in Molecular Biology 2016: 115:131.115.111-131.115.113.

17 Joung, J. et al. Genome-scale CRISPR-Cas9 knockout and transcriptional activation screening. Nat Protoc 2017; 12: 828–863.

18 Jost, M. et al. Combined CRISPRi/a-Based Chemical Genetic Screens Reveal that Rigosertib Is a Microtubule-Destabilizing Agent. Mol Cell 2017; 68: 210–223 e216.

19 Szlachta, K. et al. CRISPR knockout screening identifies combinatorial drug targets in pancreatic cancer and models cellular drug response. Nat Commun 2018; 9.

20 Kampmann, M. Elucidating drug targets and mechanisms of action by genetic screens in mammalian cells. Chem Commun 2017; 53: 7162–7167.

21 Neggers, J. E. et al. Identifying Drug-Target Selectivity of Small-Molecule CRM1/XPO1 Inhibitors by CRISPR/Cas9 Genome Editing. Chem Biol 2015; 22: 107–116.

22 Wang, M. & Kaufman, R. J. Protein misfolding in the endoplasmic reticulum as a conduit to human disease. Nature 2016; 529: 326–335.

23 Hetz, C., Zhang, K. & Kaufman, R. J. Mechanisms, regulation and functions of the unfolded protein response. Nat Rev Mol Cell Biol 2020; 21: 421–438.

24 Sano, R. & Reed, J. C. ER stress-induced cell death mechanisms. Biochim Biophys Acta 2013; 1833: 3460–3470.

25 Hetz, C. & Papa, F. R. The Unfolded Protein Response and Cell Fate Control. Mol Cell 2018; 69: 169–181.

26 Bergmann, T. J. & Molinari, M. Three branches to rule them all? UPR signalling in response to chemically versus misfolded proteins-induced ER stress. Biol Cell 2018; 110: 197–204.

27 Athanasiou, D. et al. The role of the ER stress-response protein PERK in rhodopsin retinitis pigmentosa. Hum Mol Genet 2017; 26: 4896–4905.

28 Luhr, M. et al. The kinase PERK and the transcription factor ATF4 play distinct and essential roles in autophagy resulting from tunicamycin-induced ER stress. J Biol Chem 2019; 294: 8197–8217.

29 Credle, J. J., Finer-Moore, J. S., Papa, F. R., Stroud, R. M. & Walter, P. On the mechanism of sensing unfolded protein in the endoplasmic reticulum. Proc Natl Acad Sci U S A 2005; 102: 18773–18784.

30 Zhou, J. et al. The crystal structure of human IRE1 luminal domain reveals a conserved dimerization interface required for activation of the unfolded protein response. Proc Natl Acad Sci U S A 2006; 103: 14343–14348.

31 Yoshida, H., Matsui, T., Yamamoto, A., Okada, T. & Mori, K. XBP1 mRNA is induced by ATF6 and spliced by IRE1 in response to ER stress to produce a highly active transcription factor. Cell 2001; 107: 881–891.

32 Calfon, M. et al. IRE1 couples endoplasmic reticulum load to secretory capacity by processing the XBP-1 mRNA. Nature 2002; 415: 92–96.

33 Shen, X. et al. Complementary signaling pathways regulate the unfolded protein response and are required for C. elegans development. Cell 2001; 107: 893–903.

34 Acosta-Alvear, D. et al. The unfolded protein response and endoplasmic reticulum protein targeting machineries converge on the stress sensor IRE1. Elife 2018; 7.

35 Haze, K., Yoshida, H., Yanagi, H., Yura, T. & Mori, K. Mammalian transcription factor ATF6 is synthesized as a transmembrane protein and activated by proteolysis in response to endoplasmic reticulum stress. Mol Biol Cell 1999; 10: 3787–3799.

36 Ye, J. et al. ER stress induces cleavage of membrane-bound ATF6 by the same proteases that process SREBPs. Molecular Cell 2000; 6: 1355–1364.

37 Spaan, C. N. et al. Expression of UPR effector proteins ATF6 and XBP1 reduce colorectal cancer cell proliferation and stemness by activating PERK signaling. Cell Death Dis 2019; 10: 490.

38 Kim, D., Langmead, B. & Salzberg, S. L. HISAT: a fast spliced aligner with low memory requirements. Nat Methods 2015; 12: 357–360.

39 Liao, Y., Smyth, G. K. & Shi, W. featureCounts: an efficient general purpose program for assigning sequence reads to genomic features. Bioinformatics 2014; 30: 923–930.

40 Michael Love, S. A., Wolfgang Huber. Differential analysis of count data – the DESeq2 package. 2013.

41 Yu, G. C., Wang, L. G., Han, Y. Y. & He, Q. Y. clusterProfiler: an R Package for Comparing Biological Themes Among Gene Clusters. Omics 2012; 16: 284–287.

42 Rajapaksa, G. et al. ERbeta decreases breast cancer cell survival by regulating the IRE1/XBP-1 pathway. Oncogene 2015; 34: 4130–4141.

43 Tanenbaum, M. E., Gilbert, L. A., Qi, L. S., Weissman, J. S. & Vale, R. D. A protein-tagging system for signal amplification in gene expression and fluorescence imaging. Cell 2014; 159: 635–646.

44 Zhu, X. et al. Ubiquitination of inositol-requiring enzyme 1 (IRE1) by the E3 ligase CHIP mediates the IRE1/TRAF2/JNK pathway. J Biol Chem 2014; 289: 30567–30577.

45 Li, R. et al. beta-carotene attenuates weaning-induced apoptosis via inhibition of PERK-CHOP and IRE1-JNK/p38 MAPK signalling pathways in piglet jejunum. J Anim Physiol Anim Nutr (Berl) 2020; 104: 280–290.

46 Upton, J. P. et al. IRE1alpha cleaves select microRNAs during ER stress to derepress translation of proapoptotic Caspase-2. Science 2012; 338: 818–822.

47 Lee, A. H., Iwakoshi, N. N., Anderson, K. C. & Glimcher, L. H. Proteasome inhibitors disrupt the unfolded protein response in myeloma cells. Proc Natl Acad Sci U S A 2003; 100: 9946–9951.

48 Matsumoto, H. et al. Selection of autophagy or apoptosis in cells exposed to ER-stress depends on ATF4 expression pattern with or without CHOP expression. Biol Open 2013; 2: 1084–1090.

